# Multiple instance learning on tile level-pathology images provides accurate and interpretable classification for breast cancer molecular subtypes

**DOI:** 10.64898/2025.12.19.695372

**Authors:** Konstantinos Athanasios Papagoras, Ole Lund, Carolina Barra

## Abstract

Accurate breast cancer molecular subtyping is critical for treatment decisions, yet standard methods such as immunohistochemistry and gene expression profiling are costly and labor intensive. Deep learning classification approaches using Hematoxylin and Eosin-stained whole slide images are an active area of research. However, many existing methods rely on large, high-quality annotated datasets where tumor regions are manually outlined for segmentation. This process is costly, does not scale well, depends on expert pathologists, and may ignore relevant tumor microenvironment features or reflect subjective labelling decisions. Here, we present an annotation-free, weakly supervised pipeline and web-based tool for breast cancer molecular subtyping using a computational pathology foundation model. A total of 1433 WSIs from three public cohorts (TCGA-BRCA, CPTAC-BRCA, and the Warwick HER2 cohort) were tiled into 224×224 patches without overlap at 20× magnification. Tile-level embeddings were extracted with a foundation model, and slide-level representations were obtained by mean pooling. We evaluated one-vs-rest classifiers including cosine similarity, logistic regression, and attention-based multiple instance learning. On a held-out test set of 287 WSIs, calibrated logistic regression achieved a macro F1 score of 0.75 using slide embeddings, while attention-based MIL reached 0.83 using tile embeddings. Luminal A and Basal subtypes were predicted reliably, whereas Luminal B remained challenging. Novel attention and probability heatmaps highlights spatial regions most informative for predictions, supporting qualitative interpretability. These results demonstrate accurate and interpretable breast cancer subtyping without tumor annotations, and we provide a web server to support pathology diagnostics.

## 1. Introduction

In 2025, breast cancer remains a major global health concern, accounting for approximately 12.5% of all cancer cases, with projections estimating around 3 million new diagnoses by 2040. ^1^ Breast cancer is a heterogeneous and complex disease, classified into four primary subtypes: Luminal A (LumA), Luminal B (LumB), HER2-enriched (HER2), and Basal-like (BL), each defined by distinct characteristics and biomarker expression. ^2, 3^ Differentiating between these subtypes can be challenging due to overlapping histological features, often requiring gene expression profiling and immunohistochemical (IHC) analysis. Accurate classification is clinically critical, as treatment strategies differ significantly across these subtypes. ^4^ Clinically, LumA tumors are primarily treated with endocrine therapy, ^5^ LumB tumors typically require a combination of endocrine therapy and chemotherapy, ^6^ whereas HER2-tumors necessitate HER2-targeted therapies such as trastuzumab, and tyrosine kinase inhibitors such as lapatinib in addition to chemotherapy. ^7^ Hematoxylin and eosin (H&E) staining is widely used for this task, as it provides an effective balance between time efficiency and cost. ^8^

In digital pathology, H&E-stained histopathological images have increasingly been leveraged with machine learning (ML) based predictors for cancer classification. ^9^ Within breast cancer specifically, several studies have demonstrated the potential of such approaches for molecular subtype prediction. However, key challenges remain, including subtype heterogeneity, ^10, 11^ variability in H&E sources, ^12^ limited external validation across datasets from different institutions and scanners, raising concerns about generalizability. Lastly, a lack of explainability restricts clinical integration. ^13^

Many recent studies assume access to large, finely annotated datasets or rely on tumor segmentation to guide training. While effective, these approaches reduce reproducibility and accessibility, or when working with legacy datasets lacking detailed labels. Following earlier work on this dataset, ^14^ we developed a simpler, and more resource-efficient pipeline, by utilizing UNI-2, ^15^ a pathology foundation model as a feature extractor and later used those representations to develop models in a reduced feature space.

Feature extraction transforms raw data (e.g., image pixels) into lower dimensional numeric representations that capture important information for downstream tasks. In computational pathology, this is implemented via transfer learning: deep neural networks pretrained on large datasets are used as fixed backbones or fine-tuned to produce per-image embeddings. ^16^

This study provides a breast cancer molecular subtypes classifier using only slide-level labels and compact WSI feature representations extracted from UNI-2 and investigates methods performance. The key advantage of this weakly supervised classifier relies on providing a pipeline to accurately predict cancer subtypes without the need for an extensive pathology annotation of the data used to train the model. Our goal therefore was, to combine performance, interpretability, and computational efficiency within an annotation-free framework trained on a comparatively small dataset, offering a practical and accessible approach to breast cancer subtype classification.

To explore the structure of the slide-level representations, we applied dimensionality-reduction techniques including principal component analysis (PCA) and linear discriminant analysis (LDA). We benchmarked several classification strategies: k-nearest neighbors (k-NN) using cosine similarity on mean-pooled slide embeddings; logistic regression (LR) trained directly on slide-level embeddings; and multiple instance learning (MIL) frameworks that treat tiles as instances within slide-level bags, including attention-based MIL using tile embeddings and a tile-level LR-MIL variant. Probability calibration (temperature scaling) and optional class-balancing strategies were incorporated to assess their effects on model reliability and minority-class performance. For interpretability, tile-level attention scores and spatial attention heatmaps were generated for each molecular subtype, enabling visualization of histologically relevant regions. All model implementations and experimental code used in this study are publicly available at: [https://github.com/kgoras1/BRC_ML].

## 2. Methods

Here we constructed an automated, annotation-free, and weakly supervised breast cancer molecular subtyping from H&E WSIs. The workflow consists of four main stages: (1) dataset assembly and subtype-cohort data splits; (2) WSI preprocessing and tiling; (3) tile-level feature extraction using the pathology foundation model UNI-2; and (4) model development and evaluation across multiple learning scenarios.

### 2.1 Dataset

#### 2.1.1 Breast Cancer Molecular Subtypes Cohorts

Following prior work, we collected WSI resources for breast cancer including slides with molecular subtype labels from TCGA-BRCA and CPTAC-BRCA ^17,18^ to mirror the references splits. ^14,19^ Table 1 summarizes the composition of the datasets, detailing labeling sources, molecular subtype distributions, and excluded slides. From TCGA we downloaded 1116 H&E WSIs; 971 had subtype labels (BL, HER2, LumA, LumB). From CPTAC we obtained 653 WSIs, of which 382 were subtype-labeled (BL, HER2, LumA, LumB). Slides labeled as Normal-like and those lacking labels or corrupted were excluded. Across the combined subtyped-labeled set (n=1353), LumA was the majority class (50.1%) and HER2 the minority (8.5%), indicating substantial imbalance. To partially address this, we also gathered the Warwick cohort used by the reference, ^20^ comprising 88 WSIs with 80 HER2-positive slides used in our analyses. Table 2 summarizes the technical characteristics of the breast H&E whole-slide images used in this study, including scanner type, pixel resolution, and image format.

**Table 1.** Number of WSIs per molecular subtype included in the classification dataset.

| Dataset | Labels Source | LumA | LumB | HER2 | BL | Excluded |
| --- | --- | --- | --- | --- | --- | --- |
| TCGA-BRCA | PAM50* | 502 | 221 | 76 | 172 | 35 Normal-like 110 lack labels, or corrupted |
| CPTAC-BRCA | PAM50* | 176 | 44 | 39 | 123 | 13 Normal-like and 258 lack labels |
| HER2-Warwick | IHC | - | - | 80 | - | 8 with negative HER2 expression score |
| Total | - | 678 | 265 | 195 | 295 | 424 |

\*PAM50 is a gene expression-based assay that classifies breast tumors into the four intrinsic molecular subtypes (LumA, LumB, HER2 and BL) using the expression patterns of 50 selected genes.<sup>21</sup>

**Table2.** Technical characteristics of the breast H&E slides used in this study.

| **Dataset Source** | **# WSIs** | **Pixel Size (µm/pixel)** | **Scanner Type** | **Image format** |
| --- | --- | --- | --- | --- |
| TCGA-BRCA | 971 | 0.25, 0.50 | Variant | svs |
| CPTAC-BRCA | 382 | 0.25, 0.50 | Variant | svs |
| HER2-Warwick | 80 | 0.23 | NanoZoomer C9600 | ndpi |

To ensure fair model selection and evaluation, the dataset was divided into training and test sets using an 80/20 split, with fixed random seeds for reproducibility. Stratified sampling was applied based on both molecular subtype and data source (TCGA, CPTAC, Warwick) to maintain balanced representation across splits. All classifiers were trained and evaluated on these identical splits, ensuring performance comparisons were based on completely unseen test data.

For hyperparameter tuning, model selection, and probability calibration, the training set was further divided into training, validation, and calibration subsets in an 8:1:1 ratio, maintaining stratification by class and data source. These splits were applied globally across all binary classifiers to ensure fair comparison. Sample and patient identifiers were consistently tracked to prevent data leakage. For MIL models, all tiles belonging to the same patient (i.e., from the same WSI) were grouped and assigned to a single split, ensuring that no information from a given case was shared between training and evaluation sets.

### 2.2 Pre-processing

We applied a uniform preprocessing pipeline to all cohorts. WSIs were tiled with the OpenSlide library at a nominal 20× magnification into non-overlapping 224 × 224 RGB patches; when a slide lacked an explicit 20× level the nearest available level was resampled to achieve a comparable microns-per-pixel resolution. For each tile, we saved its slide coordinates (location at the specified level) so the results can be projected back onto the original WSI for heatmaps generation and interpretation. To preserve an annotation-free, weakly supervised workflow we did not use a tumor/non-tumor detector but kept all tissue tiles.

We retained tiles containing meaningful tissue and removed those that were visually uninformative. Tiles were discarded if more than 80% of pixels exceeded a grayscale intensity of 230 (predominantly white background), or if they exhibited very low texture content were removed. Here, texture was quantified using global intensity statistics: tiles with a grayscale intensity standard deviation < 15 (indicating low contrast) or a Laplacian variance < 100 (indicating a lack of high-frequency spatial detail such as edges or structural variation). The same filtering criteria was applied consistently across TCGA-BRCA, CPTAC-BRCA, and Warwick. Tiles were stored as JPEG images. No stain normalization or data augmentation was used, ensuring that downstream performance reflects the modeling approach rather than heavy preprocessing or tumor preselection. This procedure yielded 12.534.150 tissue tiles. The preprocessing pipeline is presented in Figure1A.

**Figure 1.**
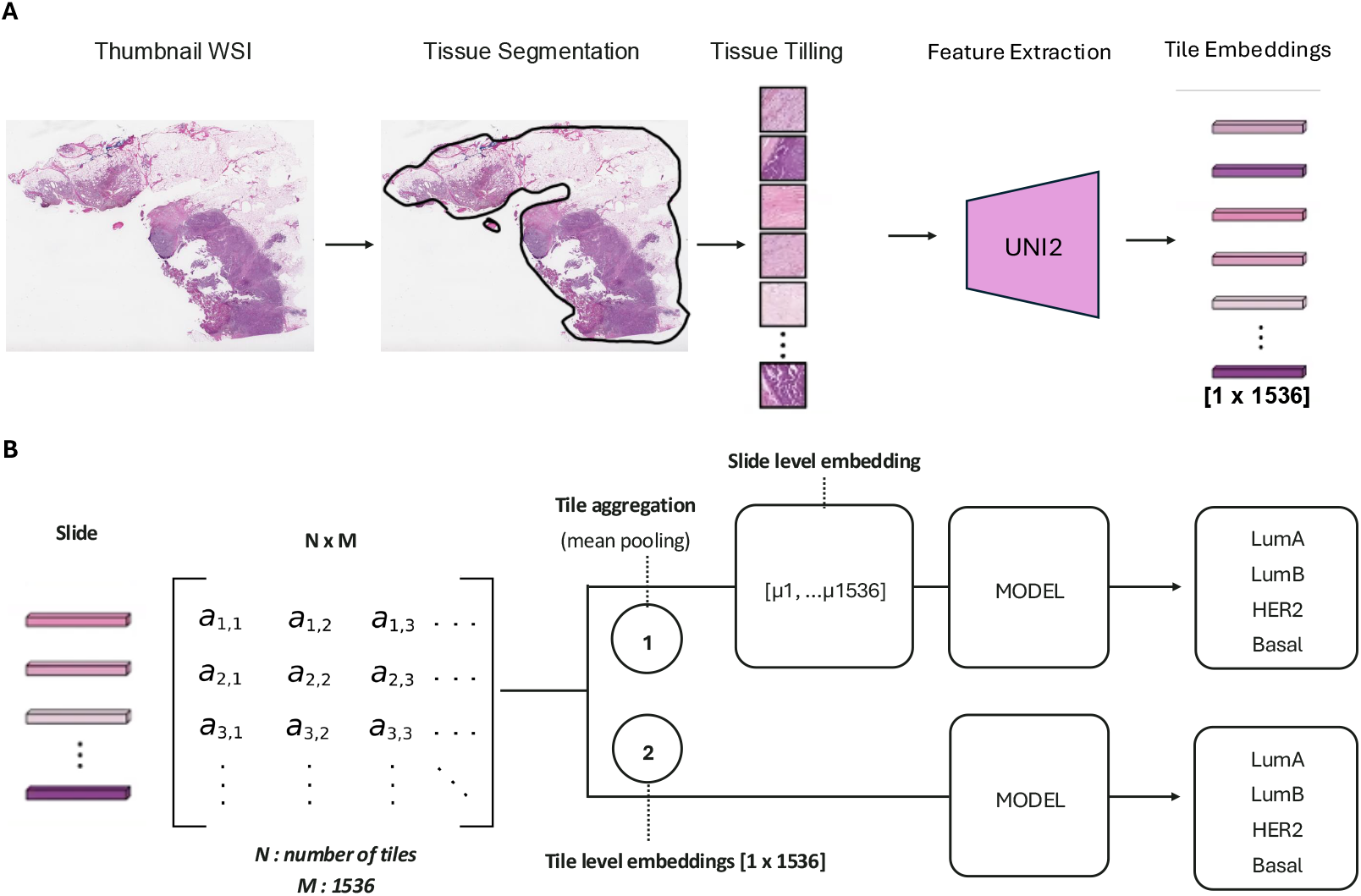
**A**. Overview of the WSI preprocessing pipeline. A WSI is first used for tissue segmentation, followed by tissue tiling. Features are extracted from each tile using the pathology foundation UNI2 model, resulting in tile embeddings (1 × 1536). **B**. Overview of slide representation as a matrix and approaches of using tile embeddings for subtype prediction. Tile embeddings are either aggregated via mean pooling (1) or grouped as instances of slide (2) and used as input for machine learning models.

This uniform strategy also mitigates site-specific bias, which is important because the Warwick cohort consists mainly of biopsy specimens and therefore yields far fewer tiles per slide compared to resection samples.

### 2.3 Feature Extraction

Following established practice and prior work such as Jaber et al. on TCGA-BRCA, ^22^ we applied transfer learning to extract tile-level embeddings from a computational pathology foundation model for breast cancer subtype classification. Based on recent reviews ^23^ and independent benchmarking studies, ^24,25^ we selected UNI-2 as the feature extractor. UNI-2 is a ViT-H/14 architecture trained with the DINOv2 self-supervised framework ^26^. Each tile is 224 x 224 x 3 pixels image, provided as a three-channel RGB image (corresponding to the red, green, and blue color channels of the H&E-stained slide). In our pipeline, UNI-2 was used as a frozen encoder, taking as input a tile image and producing a 1536-dimensional feature embedding per tile. Each slide is represented as a bag in a matrix format [number of tiles x 1536]. These slide-level bags served as the inputs to downstream subtype classification models, either through mean-pooled slide embeddings or by using full set of tile level embeddings within the MIL frameworks as illustrated in Figure 1B.

To visualize the distribution of molecular subtypes in feature space, we first generated slide-level representations by mean pooling the 1536-dimensional tile embeddings. For each slide with n tiles, the element-wise average across all tile vectors was computed, yielding a single 1536-dimensional vector per slide. These slide-level vectors were then projected to two dimensions using PCA and LDA for visualization of potential subtype clustering.

### 2.4 Classification of Breast Cancer Molecular Subtypes, Model Training

We classified breast cancer molecular subtypes using both mean-pooled slide-level embeddings and per-tile embeddings across four modeling strategies: (1) a cosine-similarity baseline, (2) LR with optional class balancing and probability calibration, (3) an attention-based deep MIL model operating on tile-level embeddings, and (4) a tile-level LR classifier. All models were trained using a one-versus-rest (OvR) scheme, where a separate binary classifier was learned for each subtype. At inference, each classifier produced a probability corresponding to its target class, and the subtype associated with the highest predicted probability was assigned as the final slide-level label.

#### 2.4.1 Cosine Similarity on Slide-level embeddings

The cosine similarity baseline predicts molecular subtypes by computing the cosine similarity between mean-pooled UNI-2 features of each test slide and all training slides. For every test slide, the *k = 3* most similar training slides are identified based on cosine similarity. Similarities are aggregated per class and transformed into class probabilities via a SoftMax normalization. The predicted subtype corresponds to the class with the highest probability.

#### 2.4.2 Logistic Regression on Slide-Level Embeddings

For the parametric LR models, we trained OvR classifiers using slide-level embeddings obtained from mean-pooled tile features. Each molecular subtype was modeled with an independent LR classifier, and hyperparameters (regularization strength and solver) were optimized via grid search with 5-fold cross-validation (cv) on the training set, using balanced accuracy as the scoring metric. Νo class balancing or probability calibration was applied in this baseline setting.

For calibrated LR, classifiers were trained and subsequently calibrated using the held-out calibration subset of the training set, following the previously described splitting strategy (2.1.1). Temperature scaling was applied to adjust predicted probabilities, which were then normalized across classes during inference to produce a single, reliable multiclass prediction per slide.

To address class imbalance, three strategies were evaluated within the same training set splits: random oversampling, random down sampling, and SMOTE. Each method was applied both with and without probability calibration, ensuring that calibration always used the designated calibration subset.

#### 2.4.3 LR on tile-level embeddings

As a classical baseline, we trained OvR LR classifiers on individual tile embeddings, using balanced class weights to mitigate class imbalance. Hyperparameters (regularization strength and solver) were optimized with the same cv method in 2.4.2 and calibrated in the calibration split via temperature scaling. Slide-level predictions were obtained by aggregating tile-level outputs using mean tile probability, and final performance was evaluated on the held-out test set.

#### 2.4.4 Attention-based Deep Multiple Instance Learning

We implemented an attention-based deep MIL framework using a OvR strategy, training a separate binary model for each class. Each binary Attention MIL model receives per-tile embeddings (1536 dimensions) and maps them to a 256-dimensional hidden representation via a linear layer, followed by ReLU activation and optional dropout. An attention head then computes a scalar score for each tile by applying a linear layer (256 to 128), followed by a tanh activation, and a second linear layer (128 to 1). The raw scores are normalized with a softmax across all tiles in the same bag to yield attention weights. The slide-level (bag) representation is computed as the attention-weighted sum of hidden tile vectors as Bag = ∑_*i*_ *a*_*i*_ *M*_*i*_ where *M*_*i*_ denotes the first linear layer, and *a*_*i*_ the attention weights. This representation is then passed to a final linear classifier (256 to 2) to produce logits for the target class versus the rest. These classifier logits were calibrated using temperature scaling on a global to all OvR unseen calibration split. Models were trained for 30 epochs using the Adam optimizer (learning rate 10^-3^), with either class-weighted cross-entropy loss or inverse-frequency oversampling to handle class imbalance. Training, validation, calibration, and evaluation were performed using the splits described in Section 2.1.1.

### 2.6 Slide-Level Prediction and Evaluation

Slide-level score computation across model types calculated as follows. For models operating on slide-level embeddings, each OvR classifier directly produced a single slide-level score per class. For Attention MIL models, slide-level scores were obtained via attention-weighted pooling of tile embeddings, yielding one score per class per slide. For tile-level LR, each tile was assigned a class probability, and the slide-level score was computed as the mean of tile probabilities within this slide.

Probability calibration was applied using temperature scaling to ensure that the output probabilities of the OvR classifiers were comparable across classes. ^27^ For multiclass prediction, each test WSI was assigned to the class with the highest calibrated probability across the four OvR classifiers. This corresponds to an argmax operation over the row-normalized calibrated probabilities, resulting in a single predicted label per slide. Importantly, no fixed per-class decision threshold (e.g., 0.5) was applied for final label assignment. The multiclass confusion matrix and overall performance metrics were computed on the test set using this calibrated argmax-based decision rule. In all cases, the resulting slide-level scores were subsequently calibrated using temperature scaling, collected across OvR classifiers, and combined via the argmax rule described above to obtain the final multiclass prediction.

For all OvR approaches using mean-pooled features, as well as the cosine similarity baseline, each binary classifier was evaluated separately using per-class metrics: accuracy, balanced accuracy, macro and weighted F1 scores, area under the receiver operating characteristic curve (ROC-AUC), and average precision (AP). Precision-recall (PR) and ROC curves were also generated. Macro and weighted averaged metrics were computed across all classes to provide an overall summary.

### 2.7 Explainability

To interpret the MIL model’s decisions, we retrieved the attention scores produced by the attention mechanism for all four subtypes for each slide. Attention heatmaps were then created to visualize the relative importance of each tile according to the model’s attention mechanism. These heatmaps highlight the spatial regions that are most influential in the slide-level predictions, providing insight into the model’s decision-making process.

### 2.7 Computational Resources

All experiments were conducted on a high-performance computing node equipped with 64 CPU cores (2.8 GHz), 1 TB of RAM, and two NVIDIA H100 NVL GPUs (each with 94 GB of VRAM).

## 3. Results

### 3.1 Exploratory Analysis of Slide-Level Embeddings

To explore the amount of discriminative information of slide-level embeddings, we first visualized the data using PCA. The first two principal components showed no clear separation between the four breast cancer subtypes, indicating substantial overlap in the embedding space (Figure 2A). The cumulative variance curve (Figure 2B) showed that 272 components were required to retain 95% of the variance, confirming that the embeddings are highly high-dimensional with limited low-dimensional separability. This highlights that subtype-related morphological variation is not captured in the top PCA dimensions.

**Figure 2.**
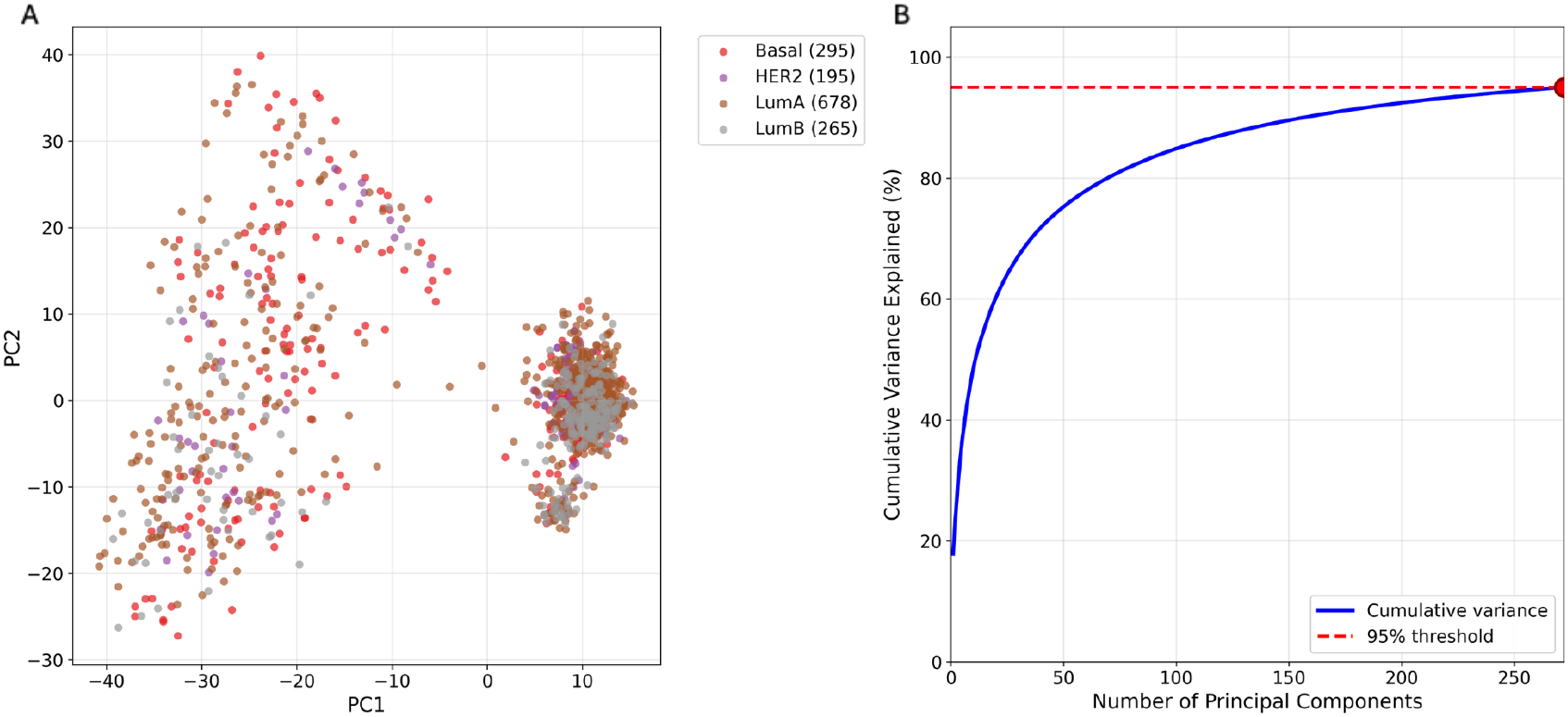
**A**. PCA scatter plot showing the first two principal components. Each point represents a slide-level embedding, colored by its molecular subtype. **B**. Cumulative variance explained by increasing numbers of principal components.

To assess whether class information could reveal structure not evident in PCA, we applied LDA, which uses subtype labels to identify projections that maximize class separability. The first two components explained 80% of the variance and revealed clear clustering of the four molecular subtypes (Figure 3).

**Figure 3.**
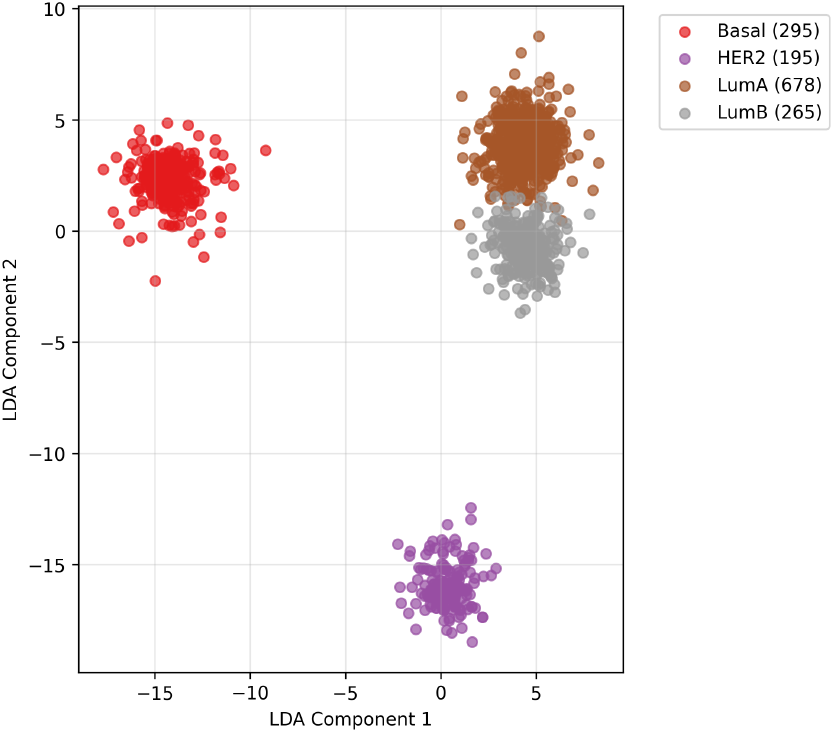
LDA projection slide-level embeddings to the first two LDA components, colored by breast cancer molecular subtype. The first two LDA components together explain 80% of the variance.

This gives a strong signal that slide-level embeddings contain discriminative information, and supervised models should be capable of subtype classification using the full embedding space.

We evaluated all models on a held-out test set of 287 WSIs (20% of the data, stratified by subtype and cohort).

To facilitate fair comparisons with prior work and between models, macro-averaged F1 scores were primarily reported, while additional evaluation metrics are presented for completeness.

### 3.2 Machine Learning on Slide-Level Embeddings

After benchmarking multiple classifiers on mean-pooled UNI-2 embeddings, we selected a calibrated OvR LR pipeline with SMOTE, which offered the best balance of performance and calibration among the evaluated models. On the held-out test set (n = 287), this approach achieved a macro F1 of 0.75. A full breakdown of precision, recall, and F1 is provided in Table 3.

**Table 3.** WSI level OvR LR (SMOTE, calibration) evaluation metrics of breast cancer molecular subtypes.

| Class | # WSIs | F1 score | Precision | Recall |
| --- | --- | --- | --- | --- |
| Luminal A | 136 | 0.84 | 0.79 | 0.89 |
| Luminal B | 53 | 0.60 | 0.68 | 0.52 |
| HER2 | 39 | 0.79 | 0.87 | 0.71 |
| Basal | 59 | 0.80 | 0.78 | 0.81 |
| Macro - average | - | 0.75 | 0.78 | 0.73 |

Performance varied substantially across subtypes. Luminal A and Basal were predicted with high reliability, and HER2 also showed strong discrimination, whereas Luminal B remained challenging, reflecting weaker separability in the embedding space (Table 3). These findings align with the confusion matrix, PR curves, and ROC curves in Figure 4, which illustrate per-class trade-offs and overall classifier discrimination. Slide-level score computation, calibration and multiclass assignment and metrics calculation are described at 2.6.

**Figure 4.**
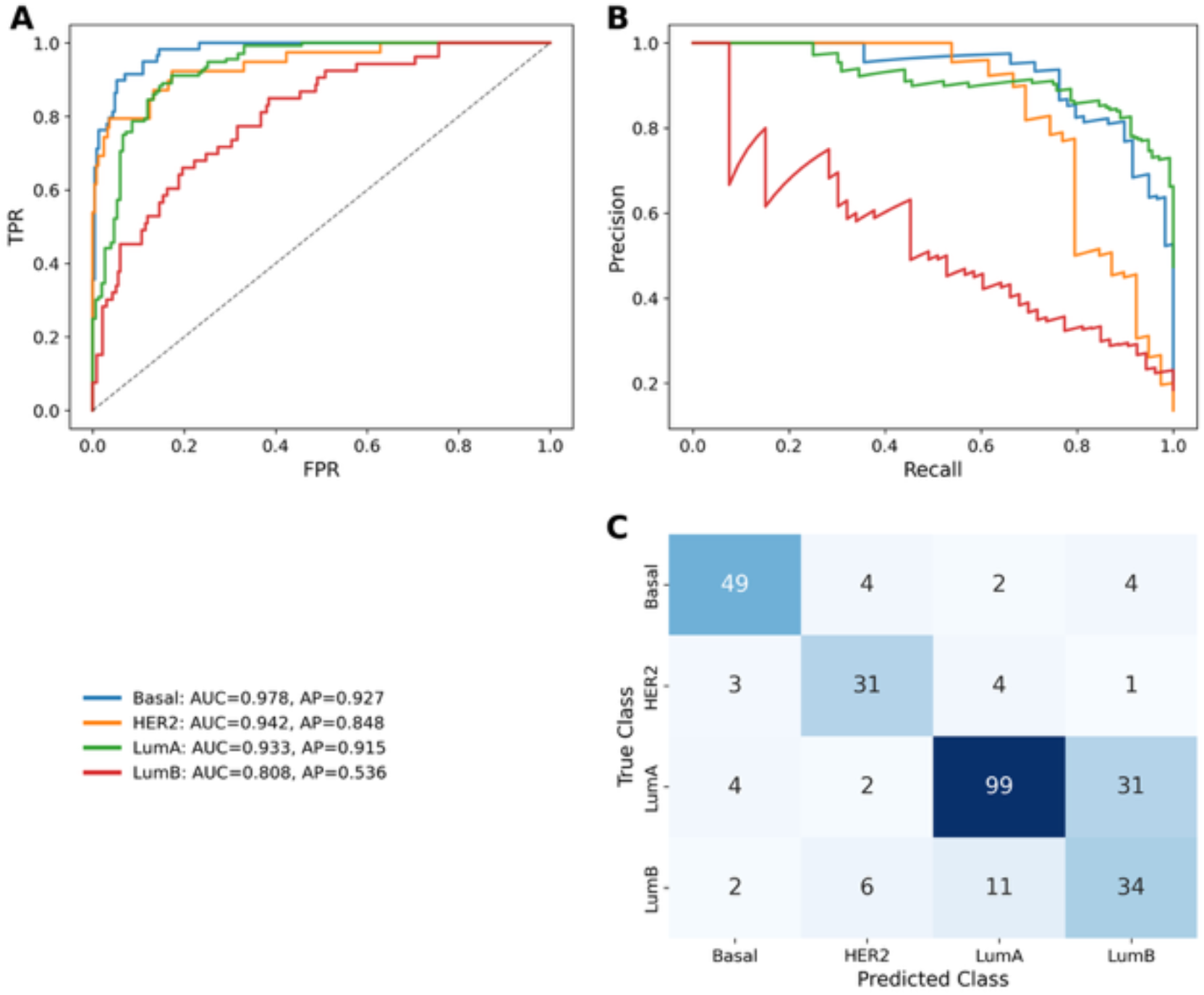
Test set performance evaluation of OvR LR (SMOTE, Calibrated) variant. **A**. PR curves of the subtypes. **B**. ROC-AUC curves of the subtypes. **C**. Confusion matrix of coupled OvR model predictions to multiclass.

The cosine similarity method provides a simple reference for evaluating model performance. From Table 4, all parametric models outperform this cosine similarity baseline. SMOTE improved class balance during training, while temperature scaling slightly reduced balanced accuracy but produced more reliable probability estimates. Performance for all competing variants including cosine k-NN, logistic regression with random oversampling, random down sampling, SMOTE, and calibrated/uncalibrated runs is summarized in Table 4. Notably, LR with SMOTE (uncalibrated) achieved the highest macro F1 (0.76), whereas applying calibration led to a small drop in macro F1 but improved probability reliability.

**Table 4.** Test set performance metrics of OvR classifiers using slide-level embeddings across different models.

| Model | Balanced Accuracy | Macro F1 |
| --- | --- | --- |
| Cosine similarity (k=3) | 0.67 | 0.68 |
| LR (Unbalanced, Uncalibrated) | 0.77 | 0.75 |
| LR (Unbalanced, Calibrated) | 0.73 | 0.75 |
| LR (Random Oversampling, Uncalibrated) | 0.77 | 0.76 |
| LR (Random Oversampling, Calibrated) | 0.73 | 0.75 |
| LR (Random Downsampling, Uncalibrated) | 0.68 | 0.64 |
| **LR (SMOTE, Uncalibrated)** | **0.78** | **0.76** |
| LR (SMOTE, Calibrated) | 0.74 | 0.75 |

### 3.2 Logistic Regression on Tile-Level Embeddings

OvR LR models trained on tile-level vectors and aggregating mean tile probabilities for slide label, achieved mean balanced accuracy 0.65, mean overall accuracy 0.73 and macro F1 0.68 on the held-out test set (n = 287). Mean precision and recall were 0.76 and 0.65, respectively. Performance was lower than Attention MIL, and training required substantially more time and memory, likely due to processing all tiles in memory without GPU acceleration.

### 3.3 Attention-based Deep MIL on Tile-Level Embeddings

The results summarize OvR Attention-MIL classifiers trained with cross entropy weighted loss function; each classifier was calibrated per class using temperature scaling on a shared calibration set, and metrics are reported as mean across classes. On the held-out test set (n = 287), the model achieved a balanced accuracy of 0.84, overall accuracy of 0.87, macro F1 of 0.83, precision of 0.72, and recall of 0.80. The full analysis of the per class metrics is shown in Table 5, along with the multiclass aggregation in the confusion matrix Figure 5 and computed as described in 2.6.

**Table 5.** Test set performance evaluation metrics of OvR Attention MIL models.

| Class | # WSIs | F1 score | Precision | Recall |
| --- | --- | --- | --- | --- |
| Luminal A | 136 | 0.86 | 0.85 | 0.86 |
| Luminal B | 53 | 0.67 | 0.40 | 0.69 |
| HER2 | 39 | 0.88 | 0.86 | 0.74 |
| Basal | 59 | 0.89 | 0.79 | 0.89 |
| Macro - average | - | 0.83 | 0.72 | 0.80 |

**Figure 5.**
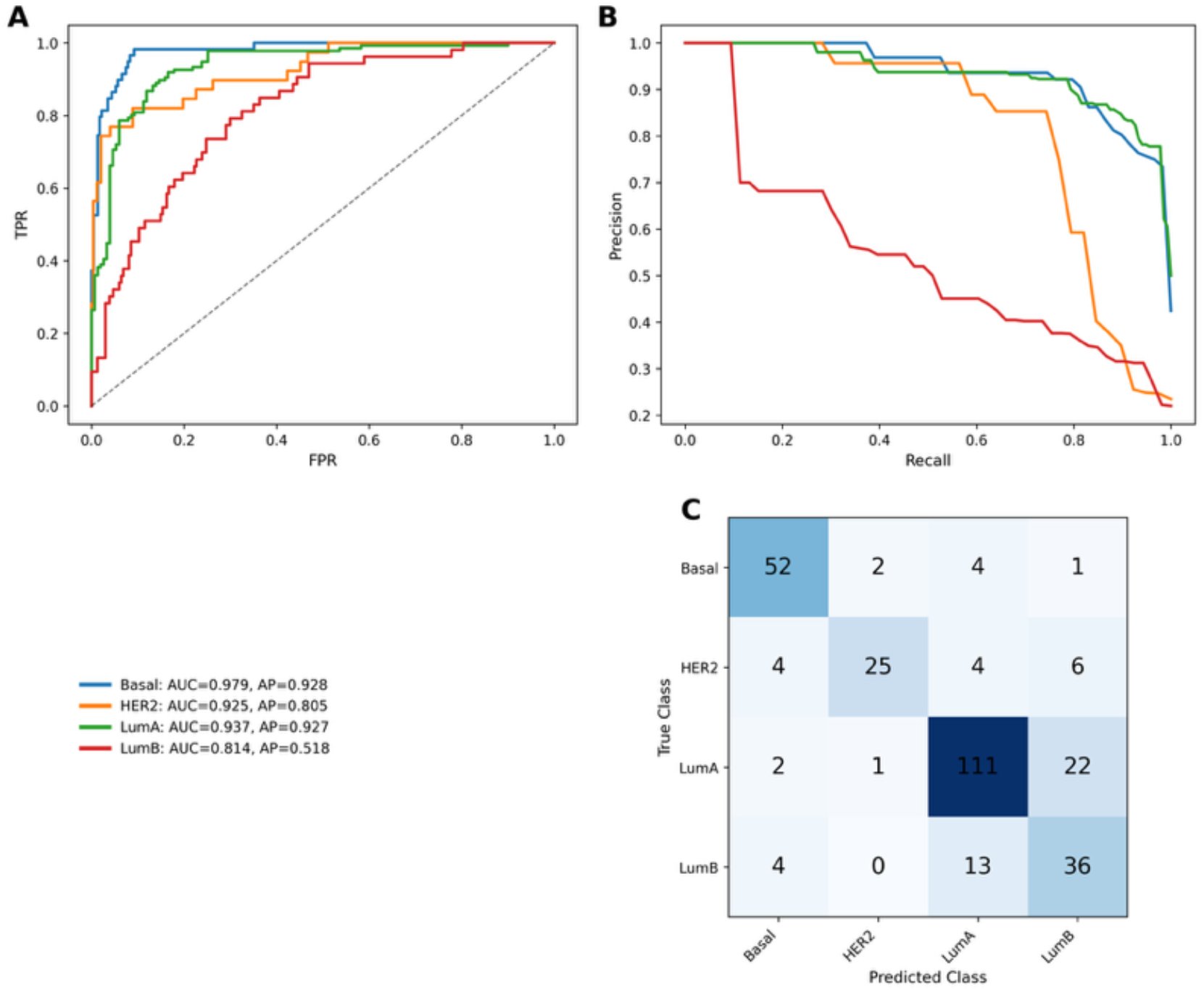
Test set performance evaluation of the OvR Attention MIL (Calibrated). **A**. PR curves of the subtypes. **B**. ROC-AUC curves of the subtypes. **C**. Confusion matrix of combined OvR model predictions to multiclass.

Notably, the calibrated Attention MIL models trained on tile embeddings achieved higher mean accuracy (0.87), mean balanced accuracy (0.84), and macro F1 (0.83) compared to the selected LR variant trained on slide-level embeddings. LumA, Basal and HER2 showed strong performance (F1 = 0.86, 0.89, 0.88) and LumB remained the most challenging subtype (F1 = 0.67).

To qualitative asses the spatial information of model’s predictions, we generated attention heatmaps for each molecular subtype classifier overlaid on H&E whole-slide images (Figure 6). These heatmaps visualize the learned attention weights that the model assigns to each tile embedding and projected on the slide during inference. Warmer colors (yellow/orange) indicate tiles with higher attention weights that drive the classification decision, while cooler colors (blue/purple) represent down weighted regions. Notably, some tumors may contain multiple molecular subtypes within the same slide, ^28^ suggesting that these heatmaps can capture intratumoral heterogeneity and localized features that binary subtype classification might miss.

**Figure 6.**
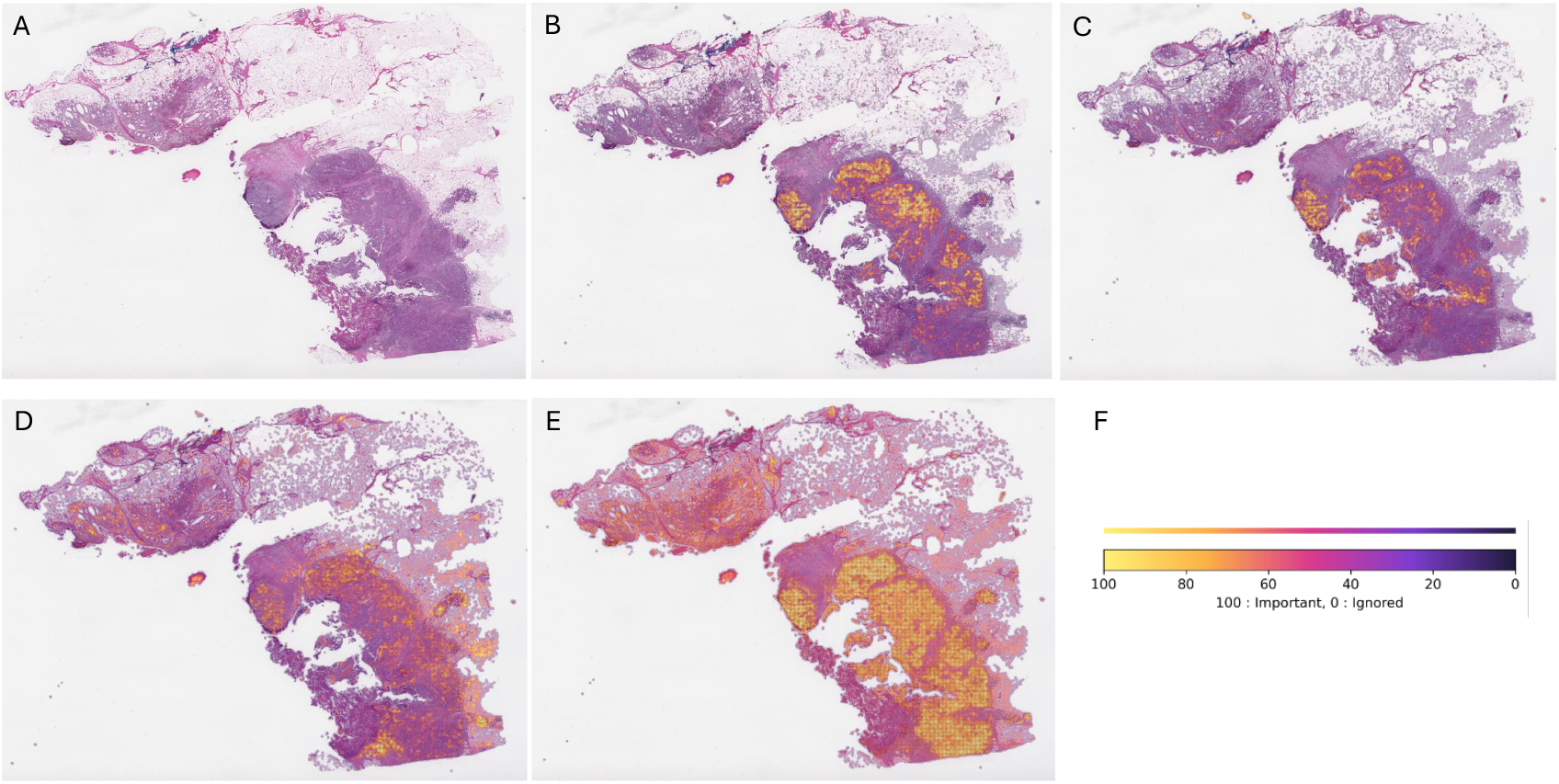
**A**. Thumbnail of the original whole-slide image. **(B–E)** Attention heatmaps generated by the Attention MIL classifiers for the four molecular subtypes: **B**. LumA, **C**. Basal, **D**. LumB, and **E**. HER2. **F**. Color bar indicating attention weights, where warmer colors denote regions of higher importance for the model’s prediction.

Across representative examples (Figure 6 B–E), the classifiers consistently focus on tumor-rich areas and each classifiers pays attention to slightly different areas. Because attention weights are learned directly during training rather than applied post-hoc, the resulting heatmaps offer an interpretable view of the spatial evidence underlying each prediction.

## 4. Discussion

In this study, we investigated whether H&E-stained WSIs contain sufficient information for molecular subtype classification using only slide-level labels, without tumor segmentation or fine-grained annotations. To address this, we utilized pathology foundation model for transfer learning, and we compared approaches based on tile-level embeddings and mean-pooled slide embeddings, evaluated with classical classifiers, such as LR, and attention-based MIL. This allowed us to assess both performance and interpretability in a resource efficient, annotation-free setting.

Across held-out tests, both slide-level and tile-level approaches recovered clinically meaningful subtype signals. The attention-based MIL model achieved the highest macro F1 score (0.83) across the OvR classifiers, outperforming the slide-level OvR logistic regression variants and the cosine-similarity baseline. In addition to improved performance, the attention mechanism enabled the generation of attention heatmaps highlighting regions contributing most strongly to subtype predictions. The slide-level OvR LR pipeline, operating on mean-pooled embeddings with SMOTE and probability calibration, also demonstrated competitive performance (macro F1 = 0.75; mean balanced accuracy = 0.74), with per-class variation consistent with the MIL model (LumA and Basal strongest, LumB weakest). Importantly, this per-subtype performance pattern was remarkably stable across all modelling approaches, indicating that the underlying separability of each subtype is driven primarily by the data rather than the specific model architecture.

Remarkably, even with this compact slide-level vector representation and weak supervision, our OvR LR approach achieved promising performance for several subtypes, highlighting its potential as an efficient pipeline for benchmarking both foundation models (as feature extractors) and ML methods (as classifiers) that can scale to larger datasets. Furthermore, the attention-based MIL model using tile level features demonstrated strong performance of weakly supervised MIL combined with foundation-model feature extraction for efficient and interpretable WSI-level breast cancer molecular subtyping. Interpretability analyses using attention heatmaps confirmed that MIL models focus on histologically relevant regions, with each binary classifier learning distinct histological patterns. These maps can serve as a form of model validation for clinicians and have the potential to guide diagnostic decisions in practice.

These trends regarding per class performance align with the reference study, ^14^ which serves as our primary benchmark, because it used the same dataset. Unlike that study, which relied on a multistage pipeline involving tissue segmentation and per-tile tumor classification to filter out non-cancerous tissue, our results show that meaningful subtype signals can be extracted directly from global WSI feature embeddings and slide-level labels. Notably, our attention-based MIL approach improves macro F1 score by 10% compared to Masoud Tafavvoghi et al., ^14^ while also providing interpretable predictions through attention heatmaps. This and other studies have explored molecular subtype prediction on different datasets, but differences in cohort composition, labeling strategies and performance metrics limit direct comparability. More specifically literature has shown that breast cancer molecular subtypes can be predicted from H&E WSIs using CNNs, self-supervised backbones, or weakly supervised MIL frameworks, but performance varies widely across datasets and evaluation settings. Reported accuracies from these studies range from ∼0.58 to 0.77 in multiclass or Basal vs non-Basal settings, with several studies highlighting challenges related to tile noise, subtype heterogeneity, and underrepresented class samples. ^29–31^ Against this, our approach achieves superior performance while providing a simpler interpretable, adjustable and computationally efficient pipeline.

Class imbalance remains a challenge particularly for HER2 and LumB. Oversampling and calibration partially mitigated this, but future work could benefit from additional data or augmentation strategies to improve minority-class performance and to understand these histological patterns that are difficult to recognize through explainable models.

Future work should validate these models on independent datasets, including potential variability in slide preparation across cohorts, explore multimodal integration (e.g., incorporating IHC or genomic data), and further develop explainable approaches to enhance transparency and support clinical adoption.

## 5. Conclusion

Overall, our results demonstrate that both tile-level and slide-level embeddings based on foundation models capture biologically relevant information for molecular subtype classification. Weakly supervised approaches, particularly attention-based MIL, provide a practical and interpretable alternative to annotation-intensive pipelines, achieving competitive performance with substantially fewer resources. While these findings highlight the promise of foundation model–based representations for breast cancer subtyping, further research could focus on optimizing performance, improving robustness and generalizability across diverse cohorts, and advancing their potential as reliable tools in clinical diagnostic workflows.

